# AE-PocketMiner Uses Attention to Simultaneously Predict Cryptic Pockets and Their Allosteric Coupling

**DOI:** 10.64898/2026.05.21.726899

**Authors:** Si Zhang, Prajna Mishra, Devin Kelly, Rachit Kumar, Gregory R. Bowman

## Abstract

Finding and targeting cryptic pockets could dramatically expand the druggable proteome. However, discovering these sites remains challenging since they are only open a fraction of the time. It is also difficult to predict the functional relevance of a cryptic site as this often requires insight into allostery. Here we introduce attention enabled (AE-)PocketMiner, an artificial intelligence (AI) method that uses a graph neural network with an attention mechanism to simultaneously predict the locations of cryptic pockets and their allosteric coupling to the rest of the protein from a single input structure. We show that AE-PocketMiner outperforms past methods for identifying cryptic pockets and recapitulates known allosteric interactions. Moreover, we experimentally confirm newly predicted cryptic pockets and mutations that allosterically control pocket opening. AE-PocketMiner thus provides a powerful framework for multiple steps of the drug discovery process—including pocket identification, prioritization, and assay design—that will help expand the druggable proteome.

## Introduction

Finding and targeting cryptic pockets is a significant challenge but success could dramatically expand the druggable proteome. These pockets are closed in the most probable structure(s) of a protein but open when the protein fluctuates to less probable conformations. Experimental methods for solving protein structures typically report on the most probable structures of a protein, leaving less probable structures that harbor cryptic pockets unresolved. A systematic way to find cryptic pockets could open a number of possibilities for drug discovery. For example, many proteins are thought to be extremely difficult drug targets, if not “undruggable,” because experimentally-derived structures of the protein lack concave pockets where small molecules are likely able to bind with the affinity and specificity required for a successful drug. Finding cryptic pockets in these proteins could provide a means to drug them^1–3^. Other classes of proteins have obvious pockets that are well-conserved across many proteins, making it difficult to specifically target one protein without also hitting the others. Finding cryptic pockets in such proteins could provide a path to achieve specificity^4–7^. Finally, drugging pockets that coincide with functional sites typically inhibits function. Targeting cryptic pockets that allosterically control distal functional sites could provide opportunities to enhance desirable functions^8–11^.

The alure of cryptic pockets has motivated the development of a number of methods for finding them. For example, a site-directed screening approach called tethering led to the development of the first FDA-approved KRAS inhibitors^1,12^. Screening fragment libraries with high-throughput crystallography has also identified cryptic pockets^13,14^. The success of these approaches depends on having sufficiently strong binders in the library of compounds that one screens. To decouple the discovery of pockets and binders, a number of simulation-based methods have been developed^15–18^. These include goal-oriented adaptive sampling strategies^19,20^ and variants of replica exchange^21,22^, among others^23,24^. Cryptic pockets identified *in silico* can then be experimentally tested using techniques like thiol labeling, hydrogen-deuterium exchange, or rational drug design^25–29^. More recently, a number of AI methods have been developed to predict cryptic pockets^30–35^. For example, PocketMiner predicts where cryptic pockets are likely to form in a single input structure based on what the model learned from training on molecular dynamics simulations^36^. A protein language model has also been developed to predict cryptic pockets based on what it learned from a large database of structures, called CryptoBank^37^. Applying these tools to large numbers of proteins suggests that cryptic pockets are quite common, highlighting the great possibilities these pockets present for drug discovery.

Despite this progress, significant hurdles to identifying and exploiting cryptic pockets remain. Even if AI can predict cryptic pockets nearly instantaneously, it is often unclear if targeting the predicted pocket would have a functional effect. It is also unclear how to use this information to screen for molecules that bind the cryptic pocket. Similarly, if one rationally designs a molecule to bind a cryptic pocket and sees that it impacts function, it can be difficult to determine whether the molecule works as intended or by a different mechanism.

To overcome these hurdles, we introduce an attention-enabled AI algorithm, called AE-PocketMiner. The attention mechanism^38^ is intended to capture long-range allosteric coupling that is beyond the local interactions encoded in the rest of the graph neural network. We assess the performance of AE-PocketMiner at scale by applying it to CryptoBank. Then we test the method on a known cryptic pocket in a membrane protein, called the Cannabinoid receptor type 1 (CB1). This is a challenging test of the model’s generalizability since our training data consists almost entirely of soluble proteins. As a next step, we experimentally test a new cryptic pocket that AE-PocketMiner predicts in δ-secretase, the enzyme responsible for cleaving the Aβ peptide that plays a prominent role in Alzheimer’s disease from its parent protein. Finally, we experimentally test AE-PocketMiner’s ability to predict mutations that modulate the probability of pocket opening in viral protein 35 (VP35). Throughout the discussion, we explain how this tool can be used to prioritize pockets for further study, to design experimental screens for molecules that bind cryptic pockets, and to test whether a proposed binder actually engages a cryptic pocket.

## Results

### Incorporating attention improves model performance

We reasoned that adding an attention mechanism, which determines which part of an input is most relevant to predicting an output, would be a valuable way of capturing allosteric coupling. In the first version of PocketMiner^36^, we represented a residue and its 30 nearest neighbors as a graph with nodes representing residues and links denoting relationships between residues. Then we updated the representation for each residue with a message passing algorithm. During training, the algorithm was presented with the initial conformation of a 40 ns molecular dynamics simulation and asked to predict the probability that each residue is part of a cryptic pocket during the snippet of simulation (i.e., the probability that a buried residue in the starting conformation becomes solvent exposed during the simulation). The model achieved excellent performance after training on 131.6 µs of simulation of 38 proteins, which corresponds to approximately 0.9 million residue-level training labels. However, message passing algorithms are limited to capturing local effects as information only passes between directly connected nodes at each iteration. While the local environment of a residue is certainly relevant for determining the probability that it is part of a cryptic pocket, there are likely long-range effects due to allosteric coupling that the first version of PocketMiner cannot capture.

To capture such allosteric effects, we added a multi-head attention module^38^ after the message passing layers (**Figure 1**). When making a prediction for residue r, this mechanism generates an attention score for every other residue in the protein that quantifies how relevant that residue is to make the prediction for residue r. Higher scoring residues have more weight in the final prediction, allowing AE-PocketMiner to capture longer range effects than a pure message passing network. If there are important allosteric effects on cryptic pocket opening, then AE-PocketMiner should outperform our previous algorithm. Examining the attention scores should also provide a facile means to determine which residues are allosterically influencing the probability of pocket opening at a given site.

**Figure 1.**
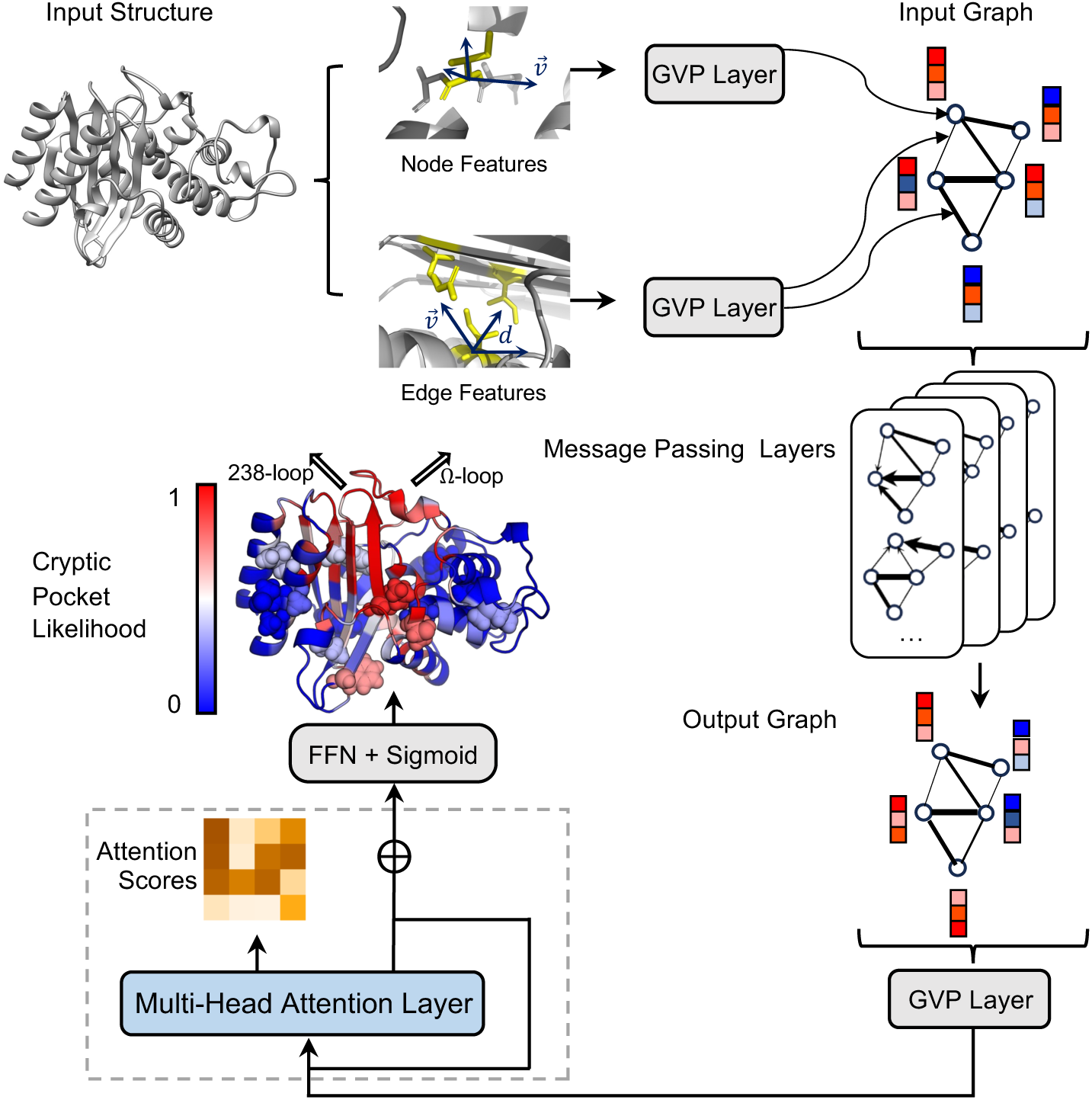
Model architecture of AE-PocketMiner. The model converts the input structure into a graph-based representation of a protein structure with nodes corresponding to residues. Nodes have features like backbone dihedral angles and edges have features encoding inter-residue relationships like radial basis functions of Cα-Cα distances (See Methods). These features are processed through geometric vector perceptron layers followed by four message passing layers that update each residue’s embeddings based on neighboring residues and connecting edges. To better capture both local and long-range residue interactions, a two-head attention block (gray dashed box) is introduced after the message passing and transformation layers. This block includes layer normalization and a residual connection that adds the original embeddings to the attended outputs. The combined embeddings are passed through a feedforward network (FFN), followed by a sigmoid activation function to predict per-residue cryptic pocket likelihood. An example prediction is shown (left middle), where residues are colored according to their predicted cryptic pocket probabilities (blue to red). The location of a known pocket is denoted with two black arrows (in the open state, the red residues in the !-loop and adjacent 238-loop shift to create the opening). The 15 residues that are predicted to have the strongest allosteric coupling to the cryptic pocket based on their attention scores are shown as spheres.

We trained AE-PocketMiner with an active learning strategy. Adding the attention mechanism increases the number of parameters that have to be learned, which tends to increase the amount of training data required. Therefore, we took an active learning approach to enlarge our training data. Specifically, we identified 86 additional proteins spanning 37 unique sequence identity clusters (see Methods) where PocketMiner mislabels residues. Then we conducted adaptive sampling simulations of these proteins and added them to our training set. The expanded training set totaled 346.4 µs of simulation time, totaling about 2.3 million residue-level data points with corresponding training labels. As described in the supporting information (**SI**; **Figure S1** and **S2**), AE-PocketMiner outperforms PocketMiner on both the validation and the final benchmark test set. These curated datasets include verified true positive and true negative cases, enabling direct evaluation of classification performance. Notably, adding attention and expanding the training data are both required to achieve this performance gain. Neither training the same architecture with attention on the original PocketMiner training set nor retrained PocketMiner on the expanded training set achieves the significant performance gains of AE-PocketMiner over PocketMiner.

We created a greatly expanded test set of 18,250 proteins from CryptoBank to test how well AE-PocketMiner generalizes. CryptoBank was created by mining the Protein Data Bank (PDB)^39^ for ligand-bound and ligand-free structures where the protein structure shifts to accommodate the ligand. Filtering out structures with missing atoms and non-natural amino acids left 744,622 pairs of ligand-bound and unbound structures. Unlike the curated test sets described above, CryptoBank does not emphasize true negatives. Residues identified as cryptic are true positives with regard to the definition of cryptic employed in CryptoBank. However, residues that aren’t labeled as cryptic could be part of as yet undiscovered cryptic pockets. Therefore, we evaluate methods based on what fraction of cryptic residues they recover, accepting that there is insufficient information to distinguish false positives and true negatives.

As expected, AE-PocketMiner significantly outperforms PocketMiner on CryptoBank. Across all residues labeled as cryptic, AE-PocketMiner achieved a recall (i.e., the fraction of true cryptic pocket residues correctly predicted) of 76.0%, substantially outperforming PocketMiner (56.7%) (**Table 1**). For high-confidence cryptic sites (mean site scores ≥ 0.5, where the mean site score is the average cryptic score across multiple holo conformations and the corresponding apo state), AE-PocketMiner recalled 72.6% of residues, compared with 47.2% for PocketMiner. These trends are consistent with ROC-AUC and PR-AUC comparisons (**Figure S2**), demonstrating that AE-PocketMiner gives improved performance and generalizes effectively to large, previously unseen datasets.

**Table 1.** AE-PocketMiner improves recovery of cryptic pocket residues on CryptoBank. Recall (fraction of annotated cryptic residues correctly predicted) is reported for PocketMiner and AE-PocketMiner on the CryptoBank-derived dataset (18,250 apo proteins; 744,622 apo-holo pairs). Residues are labeled cryptic using the CryptoBank mean site score, defined as the average cryptic score across multiple holo conformations and the corresponding apo structure, we report results for all annotated cryptic residues (mean site score ≥ 0.015) and for high-confidence site (mean site score ≥ 0.5). For each method, recall is shown using the default classification threshold (0.5) and a model-specific optimal threshold chosen by maximizing Youden’s J statistic on the validation ROC curve (PocketMiner: 0.53; AE-PocketMiner: 0.64).

| Mean Site-Score Threshold | PocketMiner |  | AE-PocketMiner |  |
| --- | --- | --- | --- | --- |
|  | Default (0.5) | Optimal | Default (0.5) | Optimal |
| 0.015 (all annotated) | 56.7% | 52.9% | 76.0% | 56.5% |
| 0.5 (high-confidence only) | 47.2% | 43.2% | 72.6% | 51.4% |

### AE-PocketMiner generalizes well beyond its training set

We tested AE-PocketMiner on the membrane protein CB1 to assess the generalizability of the algorithm beyond its main training set. Of the 124 proteins in our training set, the vast majority are soluble proteins (123 of them) and only one is a membrane protein. That membrane protein is cholesterol 25-hydroxylase (CH25H), a membrane embedded enzyme that has no structural homology to CB1. Therefore, it is interesting to assess how well-AE-PocketMiner performs on another membrane protein. CB1 (**Figure 2a**) is biologically interesting as it is a G protein-coupled receptor (GPCR) that allosterically transmits information across cell membranes. GPCRs are of great interest as therapeutic targets; there are 516 approved drugs targeting them, making up approximately 36% of all FDA-approved drugs^40^. CB1 is one of the most abundant GPCRs in the brain, where it plays roles in processes from memory to appetite and pain perception. It is a non-opioid target for pain relief, so there is great interest in drugging it to reduce pain without contributing to the opioid overdose epidemic. A recent study used computer simulations to discover a cryptic pocket in CB1and then designed a small molecule called VIP36 to target this pocket^25^. This cryptic pocket and the highly-allosteric nature of GPCRs make CB1 an excellent test case for AE-PocketMiner.

**Figure 2.**
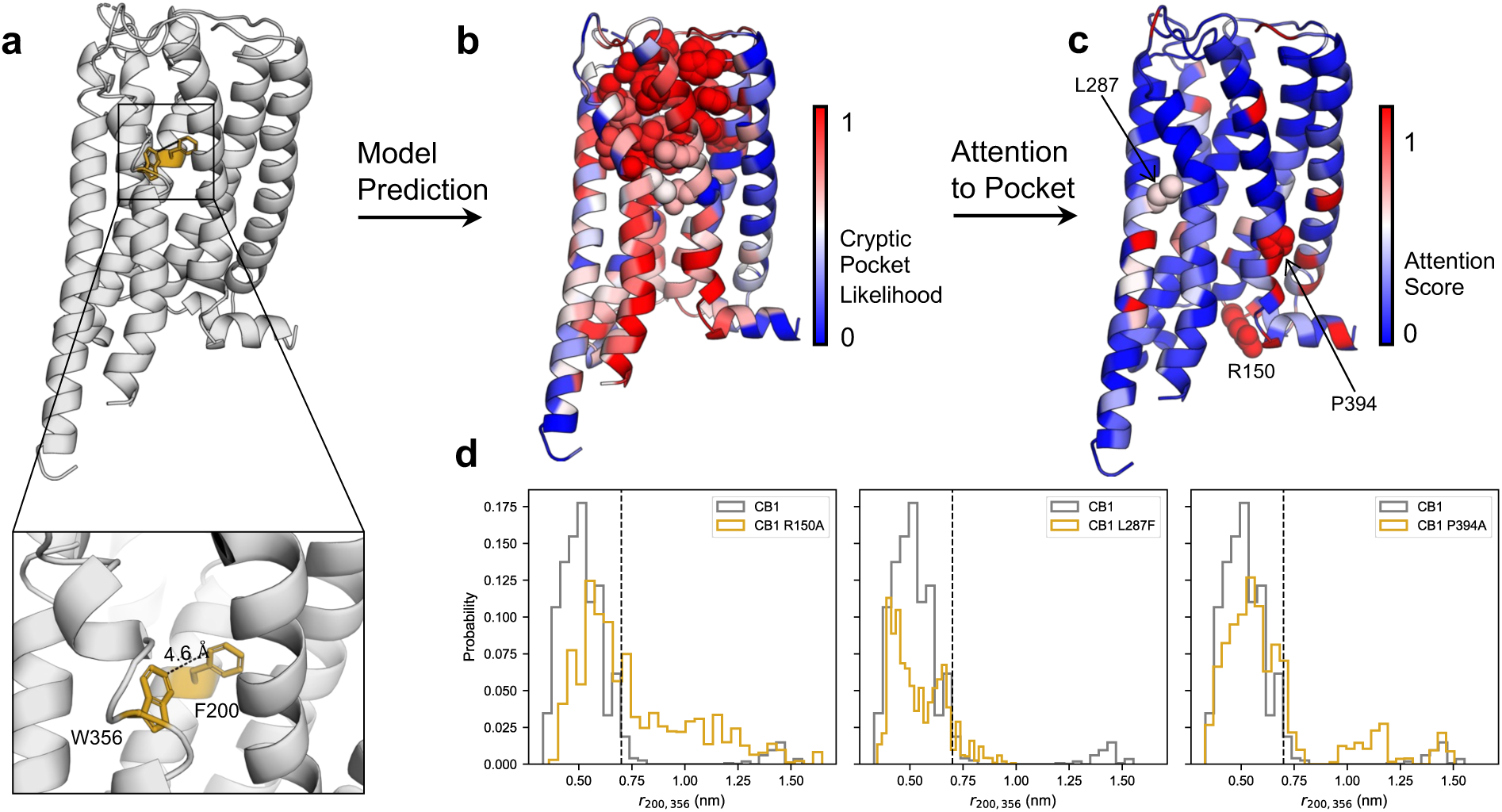
AE-PocketMiner recapitulates a pocket and allostery in the membrane protein CB1. **a)** Crystal structure of CB1 where the cryptic pocket is closed (PDB 6N4B^41^ with the FUB ligand removed). The toggle switch residues F200 and W356 are highlighted in gold; the inset shows a zoomed view highlighting the distance between CD2 of F200 and CH2 of W356. **b)** The same crystal structure with residues colored by the probability that AE-PocketMiner predicts they are part of a cryptic pocket. Residues known to line the cryptic pocket (i.e. within 5 Å of the ligand VIP36 bound in the open cryptic pocket of PDB 9B54^25^) are shown as spheres. **c)** The same structure colored by the attention scores with respect to the cryptic pocket predicted for each residue. Three residues with high attention scores-R150, L287, and P394-were selected for *in silico* mutagenesis and are shown as spheres. **d)** The allosteric effect of the selected mutations (R150A, L287F, P394A) on cryptic pocket opening, as measured by the distribution of the distance between F200 and W356 for each variant compared to wild-type (WT). The black dashed line in each plot marks the F200-W356 distance in the open crystal structure (PDB 9B54^25^; 7 Å). Larger distances correspond to pocket opening.

AE-PocketMiner does an excellent job of recapitulating the cryptic pocket in CB1 (**Figure 2b**). We defined 30 residues as being part of the cryptic pocket as they are within 5 Å of the ligand in a bound structure. AE-PocketMiner confidently predicted that 29 of these residues are part of a cryptic pocket, assigning them probabilities greater than 0.5 (**Figure S3**). It was only less confident about one residue, S390, which is on the periphery of the pocket. PocketMiner missed two additional residues. Interestingly, AE-PocketMiner also suggests there is another cryptic pocket on the opposite side of CB1. For the purposes of this work, we will focus on the known cryptic pocket.

More importantly, AE-PocketMiner provides a facile means to determine what distal residues in the protein are likely to allosterically influence the probability that the cryptic pocket opens. **Figure 2c** shows the total attention that the cryptic pocket residues pay to every other residue in the protein. Residues that are adjacent to pocket residues in the primary sequence are excluded (colored blue) as they get far more attention due to proximity and swamp out signal from distal residues. Some residues that receive high attention scores are relatively nearby, such as P394, while others are quite distal, such as R150 on the opposite end of the protein. Similar information is not available from PocketMiner, which is one of the big potential advantages of incorporating attention into AE-PocketMiner.

We chose to test whether mutations at three representative sites allosterically impact the probability of cryptic pocket opening in molecular dynamics simulations. The rationale for selecting each residue and the results are described below. For each mutation, we ran goal-oriented adaptive sampling simulations with our FAST algorithm to promote pocket opening and compared the results to wild-type (WT) CB1. Pocket opening was characterized in terms of the distribution of distances between residues F200 and W356, which lie on opposite sides of the cryptic pocket and move apart in the bound structure.

Simulations of the P394A variant enhance pocket opening (**Figure 2d**). This mutation was chosen because it is relatively nearby the cryptic pocket and has a high attention score. Literature review showed that P394 is part of the conserved NPxxY motif in transmembrane helix 7 (TM7) and has been implicated in stabilizing active conformations through a hydrogen bond network^42–44^. We hypothesized that P394 modulates pocket dynamics by regulating local flexibility in TM7. Indeed, mutation to alanine (P394A) increased the backbone RMSD of TM7 relative to WT (**Figure S4**). It also increased the probability of open pocket states, supporting our prediction that it allosterically influences opening of the cryptic pocket.

In contrast, L287F suppresses pocket opening (**Figure 2d**). This residue is similarly proximal to the cryptic pocket as P394. It received a relatively lower attention score the P394 but still stands out as more likely to be allosterically coupled to the cryptic pocket than most other residues (**Figure 2c**). This position is also noteworthy because it is a phenylalanine in CB2, a related GPCR that is not sensitive to the VIP36 ligand that binds the cryptic pocket in CB1. Therefore, we reasoned that L287F would likely reduce the probability of pocket opening in simulations. Indeed simulations of L287F had a lower probability of pocket opening than wild-type.

Finally, introducing an alanine at the highly distal residue R150 also enhances pocket opening (**Figure 2d**). This residue was selected because it received the highest attention score despite being one of the furthest residues from the cryptic pocket. We had no prior reason to suspect this residue is important based on our literature review on GPCRs. It is also surface exposed. Taken together, it is not intuitively obvious that mutating R150 should impact the probability of cryptic pocket opening, making it an interesting test of AE-PocketMiner’s predictive power. Excitingly, the R150A mutation significantly enhanced the probability of cryptic pocket opening.

Together, these results suggest that AE-PocketMiner is a significant advance over our past work. It is somewhat better at identifying a cryptic pocket in a membrane protein that is quite different from the predominantly soluble proteins in the training set. Moreover, AE-PocketMiner provides insight into allostery that can’t be obtained from PocketMiner and the importance of residues with high attention scores is confirmed in simulations. The results for R150A are particularly remarkable given how far it is from the cryptic pocket.

### Experiments confirm a newly predicted pocket in δ-secretase

We next sought to test if AE-PocketMiner can predict new cryptic pockets. To identify a suitable protein system, we screened all human proteins deposited in the PDB using the following criteria: proteins with fewer than 300 residues and containing between one and three buried unpaired cysteines. Requiring the buried cysteine allows us to use a thiol labeling assay to experimentally test if the buried cysteine is reachable by a chemical labeling reagent, which would support the existence of the cryptic pocket, as we have demonstrated in prior works^26,28^. This filtering yielded a candidate pool of approximately 10,000 proteins. To narrow the scope, we chose to focus on proteins that are implicated in Alzheimer’s disease given the huge unmet medical need that this disease presents. We further required that the buried cysteines show high attention scores relative to known functional sites, indicative of allosteric coupling, and that the protein be commercially available to expedite experimental test.

We found that δ-secretase satisfies all our criteria, particularly having two buried cysteine that AE-PocketMiner predicts are part of separate cryptic pockets, one of which has substantial allosteric coupling to the active site. This enzyme is a lysosomal protease that cleaves amyloid precursor protein (APP) to create the Aβ peptide that is well known in Alzheimer’s pathology^45^. AE-PocketMiner confidently predicts that both buried cysteines (C219 and C50) are part of cryptic pockets (probabilities of 0.59 and 0.57, respectively). C219 also has a high attention score relative to the active site (**Figure 3b**). There is one other cysteine in the protein, the catalytic cysteine C189, which is partially surface exposed. The protein is also available commercially.

**Figure 3.**
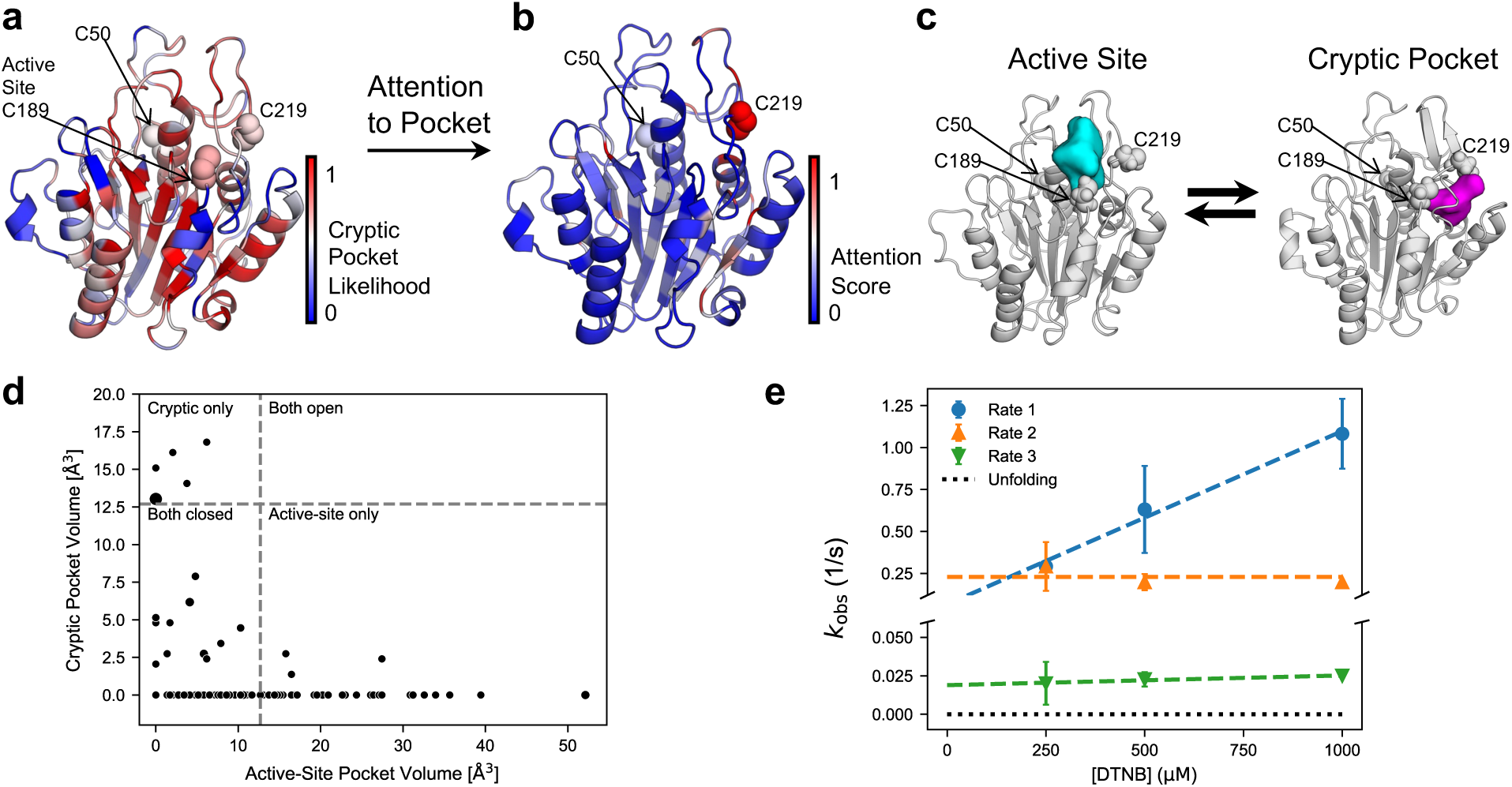
AE-PocketMiner predicts a cryptic pocket in δ-secretase that is supported by our thiol labeling experiments. **a)** Crystal structure of δ-secretase colored by AE-PocketMiner predicted cryptic pocket probabilities (PDB 5LUA^45^ with ligand 5KN removed). The catalytic cysteine (C189) and two buried cysteines (C50 and C219) are shown as spheres. Both buried cysteines have a reasonable probability of being part of a cryptic pocket. **b)** The same structure colored by AE-PocketMiner attention scores with respect to the active-site pocket. The buried C219 (highlighted as spheres) has strong coupling to the active site. **c) Representative structures when the cryptic pocket adjacent to C219 is closed (left) or open (right) suggest that opening of this pocket is anti-correlated with opening of the active site**. The cyan and magenta surfaces represent the open pocket volume in each structure and C50, C189, and C219 are shown as spheres. **d)** Scatter plot comparing the volume of the cryptic pocket adjacent to C219 to the volume of the active site for each state in our MSM supports the anti-correlation between the opening of the two pockets. For illustration, we considered either pocket to be open if its volume exceeded that of the active site in the inhibitor-bound structure 5LUA^45^ (12.69 Å^3^). Point size is proportional to the population of the state. **e)** Experimentally observed thiol labeling rates for the three cysteines (C50, C189, and C219) in δ-secretase across a range of DTNB concentrations corroborate there is a pocket adjacent to each cysteine, consistent with the cryptic pocket probabilities predicted by AE-PocketMiner. Markers show mean rates from three replicates, with error bars indicating the standard deviation. Dashed colored lines indicate linear fits to the data, and black dotted lines denote the expected labeling rate for the unfolded state.

Molecular dynamics simulations support the prediction of a cryptic pocket adjacent to C219. As shown in **Figure 3c**, our goal oriented adaptive sampling algorithm identifies a cryptic pocket that exposes the side chain of C219. The probability this pocket is open is about 0.55, in reasonable agreement with the predicted probability from AE-PocketMiner. The active site where C189 resides also fluctuates between open and closed states (open probability of 0.57, again in good agreement with the AE-PocketMiner prediction). We also find that the volume of the active site and cryptic pocket are anticorrelated (**Figure 3d**). When one pocket is open, the other pocket is likely closed, consistent with the allosteric coupling suggested by AE-PocketMiner. In contrast to the AE-PocketMiner prediction, C50 remains buried throughout our simulations. Taken together, we expect to see labeling of at least two cysteines (C189 and C219) in our thiol labeling assays, and possibly of a third (C50).

Thiol labeling experiments show labeling of all three cysteines, consistent with the AE-PocketMiner prediction that there are cryptic pockets adjacent to C50 and C219 (**Figure 3e**). All three measured rates are much faster than the expected labeling rate from the unfolded state, indicating that labeling arises from native-state fluctuations rather than unfolding. If either of our predicted pockets didn’t exist, then the adjacent cysteine shouldn’t label and we should have observed fewer than three labeling rates. While we can’t be certain which rates correspond with which cysteines, we suspect the fastest labeling rate corresponds to the active site C189 since that residue is partially exposed in the crystal structure. In light of the simulation results, we suspect the second fastest rate corresponds to the cryptic pocket by C219 and that the slowest rate corresponds to labeling of C50. All three rates are consistent with those we’ve observed for cysteines in other cryptic pockets. If any of the cysteines were predominantly surface exposed, then we would expect that cysteine to label significantly faster than any of the rates we observe. Therefore, our results corroborate that the active site is fluctuating between open and closed states. Moreover, the observation of three rates confirms there are cryptic pockets adjacent to C219 and C50.

The proteomics literature provides further experimental support for the cryptic pocket by C219 and its allosteric coupling to the active site^46^. Motivated by our success by thiol labeling, we did a deeper literature review on δ-secretase, particularly C219 since we predict this residue is part of a cryptic pocket with allosteric coupling to the active site. A study using one of δ-secretase’s alternative names (legumain) showed that one can oxidize C219 with the compound iodoacetamide-alkyne (IA). The paper did not go into the structural mechanism for labeling. However, given our observation that C219 is buried in available crystal structures, the covalent modification of C219 with IA corroborates our computational prediction of an adjacent cryptic pocket. The proteomics paper also shows that oxidation of C219 and the mutations C219S and C219A all inhibit autoproteolytic cleavage of the propeptide that must be removed for activation of δ-secretase. This observation nicely corroborates our prediction that the cryptic pocket by C219 is allosterically coupled to the active site.

Our results with δ-secretase demonstrate how AE-PocketMiner can be used to find novel cryptic pockets and allostery. Our choice of this protein shows how one can use AE-PocketMiner to choose which of a number of candidates to pursue for drug discovery. Similarly, our use of the attention scores to prioritize the pocket by C219 also shows how the tool can aid in prioritizing potential cryptic sites within a single target protein.

### Experiments confirm allosteric control over a cryptic pocket in VP35

We also reasoned that insight into allostery from AE-PocketMiner’s attention scores could allow us to design mutations that allosterically enhance or suppress the probability of pocket opening. Designing such variants could be useful for informing the design of experimental screens for molecules that target cryptic pockets. For example, one could do positive selections against a protein with a high probability of pocket opening, where the energetic cost for a hit to stabilize the open state of the pocket is reduced, and then do negative selections with variants that suppress pocket opening, as molecules that still hit them are likely to bind outside the cryptic pocket. Similarly, one could use such variants to build confidence that a molecule rationally designed to target a cryptic pocket actually binds there before committing to solve a structure. That is, a molecule that binds a cryptic pocket should be more potent against protein variants with an enhanced probability of pocket opening and less potent against variants with a reduced probability of pocket opening.

We tested whether AE-PocketMiner can inform the design of variants that control the probability of a known cryptic pocket in the interferon inhibitory domain of VP35 from Zaire ebolavirus. We previously used a combination of computer simulations and biophysical experiments to discover a cryptic pocket in VP35 that allosterically controls a critical interaction with double stranded RNA (dsRNA)^26^. This pocket forms when helix 5 (residues 305-309) moves away from the four-helix bundle, exposing residues G236, P304, A306, C307, and S310 (**Figure S5**). Several nearby residues become partially exposed as well, including T237, A238, L311, F328, L338. The open and closed states can be distinguished clearly from the distance between residues G236 and A306 in computer simulations and from thiol labeling of C307 in experiments^48^. We later identified mutations that shift the cryptic pocket opening equilibrium, including F239A (increased opening) and A291P (reduced opening). Our familiarity with the protein presents a good opportunity to test if AE-PocketMiner recapitulates what we already know and to experimentally test new predictions from the algorithm.

AE-PocketMiner nicely recapitulates the cryptic pocket and much of the allostery in VP35 (**Figure 4a-b**, **Figure S6**). All five of the residues that define the cryptic pocket are confidently predicted as belonging to a cryptic pocket (probability greater than 0.5). In contrast, the original PocketMiner algorithm only confidently predicts that one of these residues is part of a cryptic pocket. This result is consistent with our finding that AE-PocketMiner better captures the allosteric effect of distal residues. The algorithm also assigns a relatively high attention score to F239, where mutations to alanine increase the probability of pocket opening (**Figure 4c**). It assigns a low score to A291, where introducing a proline is known to reduce the probability of pocket opening.

**Figure 4.**
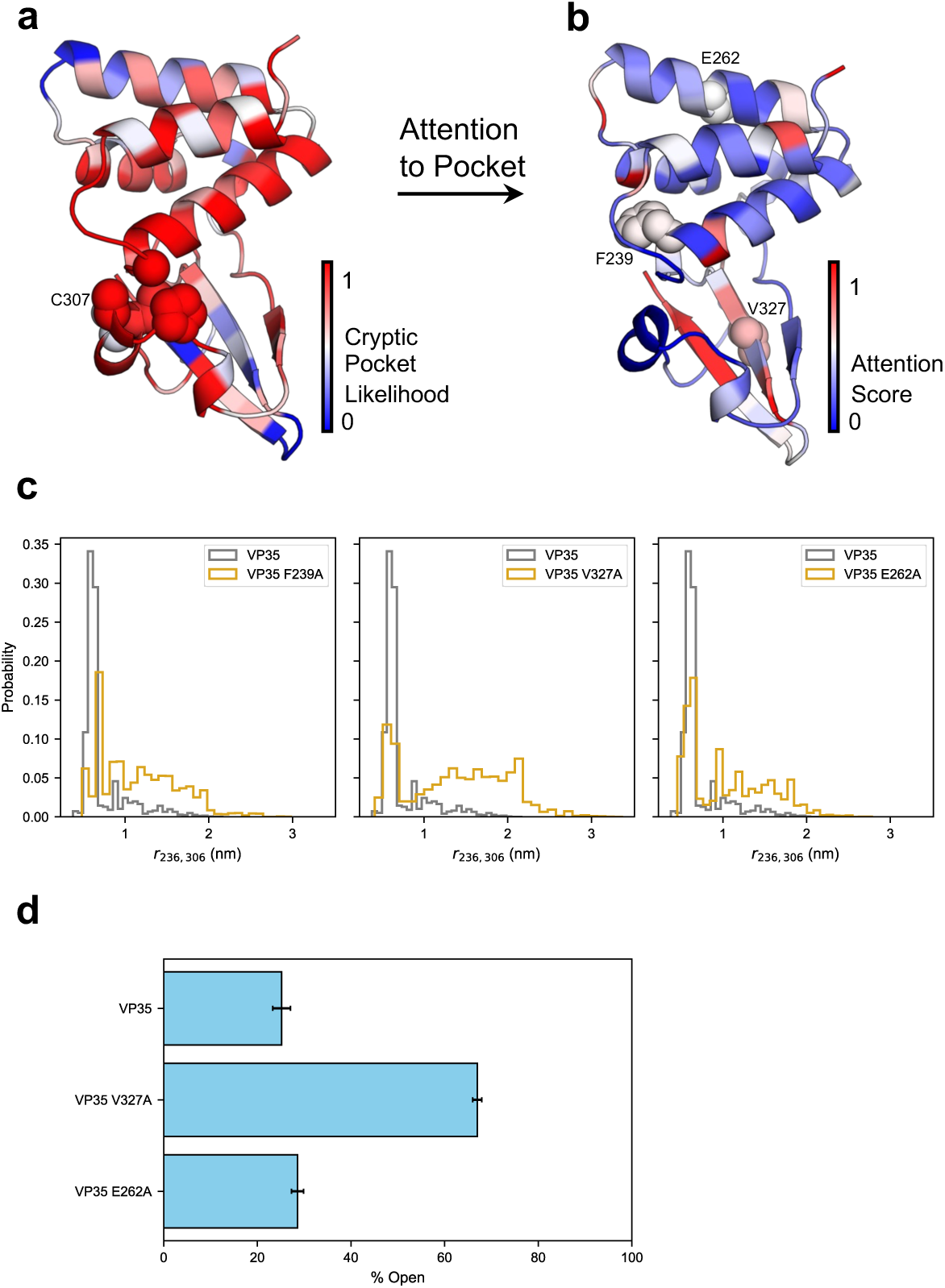
Experimental tests confirm allosteric control residues predicted by AE-PocketMiner in VP35. **a)** Crystal structure of VP35 (PDB 3FKE^47^) colored by AE-PocketMiner predicted cryptic pocket probability. Residues defining the cryptic pocket (G236, P304, A306, C307, and S310) are shown as spheres, based on prior combined experimental and computational analyses. **b)** The same structure colored by AE-PocketMiner attention scores with respect to the cryptic pocket. F239, V327, and E262 are highlighted as spheres. **c)** The allosteric effect of the selected mutations (F239A, V327A, and E262A) on cryptic pocket opening, as measured by the distribution of the distance between G236 and A306 for each variant compared to wild-type (WT). Larger distances correspond to pocket opening. **d)** The population of the open state for WT, V327A, and E262A measured by thiol labeling experiments (n=3). Error bars indicate the standard deviation across replicates.

Simulations and thiol labeling experiments also confirm that two new residues predicted by AE-PocketMiner exert allosteric control over cryptic pocket opening. We chose to characterize V327A because residue V327 receives a high attention score. It is close to the pocket (on the β-strand at the back of the pocket) but is a somewhat surprising prediction given that the sidechain points away from the pocket. We also chose to characterize E262A. This residue received a slightly lower attention score but interested us because it is very distant from the cryptic pocket and points into solution. Therefore, E262A is not an obvious choice apart from AE-PocketMiner’s prediction. Encouragingly, simulations of both variants suggest they increase the probability of pocket opening. To verify this, we conducted thiol labeling experiments and observed that V327A labels substantially faster, while E262A exhibits a modest increase in labeling relative to WT VP35 (**Figure S7**). To determine whether the enhanced labeling observed for WT, V327A, and E262A could instead arise from processes such as global unfolding, we assessed thermodynamic stability (**Figure S8**), global secondary structure (**Figure S9**), the unfolded-state population under native conditions, and intrinsic cysteine labeling rate. As shown in **Figure S10**, the observed labeling rates are significantly faster than the predicted rates for the unfolded state across of DTNB concentrations. This indicates that cysteine labeling arises from native-state fluctuations rather than global unfolding. Importantly, thiol labeling experiments are consistent with computational predictions, yielding open-state populations of 67.0 ± 1.0% for V327A and 28.6 ± 1.9% for E262A, compared with 25.2 ± 1.3% for WT (**Figure 4d**).

## Discussion

We introduced AE-PocketMiner, an attention-enabled deep learning framework that integrates a multi-head attention mechanism and a substantially expanded training dataset to enable simultaneous prediction of which residues are likely to form cryptic pockets and allosteric coupling between all pairs of residues. By incorporating attention, the model captures long-range interactions across the protein and provides a practical way to nominate distal regulatory sites for follow-up.

A key strength of AE-PocketMiner is its improved generalizability across diverse systems. AE-PocketMiner achieves higher performance than past methods on both curated benchmark test sets and the large CryptoBank evaluation. Despite limited representation of membrane proteins in the training data, the model correctly predicts the cryptic pocket in the GPCR CB1 and highlights residues that were experimentally known to allosterically modulate pocket opening.

Our experimental tests confirm the predictive power of the algorithm. For δ-secretase, a protein without any known cryptic pockets, the model identifies a previously unrecognized cryptic pocket that is functionally coupled to the active site. Subsequent simulations and thiol labeling experiments support pocket opening in this region and existing literature supports the allosteric coupling. In VP35, a system with a well-established cryptic pocket, AE-PocketMiner recapitulates the known pocket location and recovers previously characterized allosteric relationships. Moreover, it identifies additional residues where mutations are likely to allosterically modulate the probability of pocket opening, and these predictions are confirmed by our thiol labeling experiments.

We expect AE-PocketMiner to be useful for a variety of applications from obtaining mechanistic insight to protein and drug design. For example, the fact that it takes seconds to run on a given input structure means that large numbers of potential targets can be screened and prioritized based on the probability of pocket formation, the strength of allosteric coupling to key functional sites, and other criteria of interest to the user. The attention scores also provide a basis for designing mutations that allosterically enhance or suppress the probability of pocket opening. These predictions can help provide biophysical insight into how complex systems work or provide a means to design reagents for screening for molecules that target a cryptic pocket.

## Methods

### Molecular simulations

#### System preparation

The protein structures used for simulations were obtained from the Protein Data Bank (PDB)^39^. For one of the newly added proteins, ch25h, a membrane protein, the starting structure was downloaded from the AlphaFold Protein Structure Database^32,49^. SWISS-MODEL^50^ or Modeller^51^ was used to model missing loops or unresolved regions in structures, and PyMOL^52^ was employed to mutate proteins to get single– or double-point mutations.

Each protein structure was solvated in a rhombic dodecahedral box filled with TIP3P water molecules^53^, maintaining a minimum distance of 1.0 nm between the protein and the box edges. Topologies were generated using the Amber03 force field^54^ using GROMACS^55^. Na^+^ and Cl^-^counterions were added to neutralize the system at a salt concentration of 0.1 mol/liter. Protein energy minimization was performed using the steepest descents algorithm with an initial step size of 0.01 nm. Minimization continued until the maximum force dropped below 100 kJ/(mol*nm) or until 500 steps were completed. Each system was then equilibrated for 0.1 ns using a 2 fs time step. All bonds were constrained using the LINCS algorithm^56^ and long-range electrostatic interactions were calculated using the Particle Mesh Ewald (PME) method^57^ with a Fourier spacing of 0.12 nm. A cutoff of 0.9 nm was applied for both Coulomb and van der Waals interactions. The velocity-rescaling thermostat^58^ maintained the temperature at 300 K. During both equilibration and production runs, the leap-frog algorithm was used for integrating Newton’s equations of motions.

#### Adaptive sampling simulations

Adaptive sampling simulations were carried out using the FAST algorithm^19^, which iteratively guides unbiased molecular dynamics simulations toward conformations that maximize or minimize a chosen order parameter. In the first round, n parallel trajectories were launched from a same equilibrated structure. Afterward, all trajectory frames were clustered by RMSD, ranked based on the order parameter, and the top n clusters were selected to seed the next round.

For constructing the original training set, the chosen order parameter was the total LIGSITE^59^ pocket volume, which does not require prior knowledge of pocket locations. Our previous study has shown that most proteins exhibited pocket opening in the first round of FAST simulations^36^, enabling the derivation of reliable training labels from these trajectories. Therefore, we reasoned that using alternative order parameters, such as residue distances or pocket RMSD, could also be used to generate valid training labels for the newly added proteins in the expanded dataset and may even perform better when prior knowledge of the proteins is available. Trajectory frames were further clustered using the k-centers algorithm with backbone α-carbon RMSD as the distance metric. Depending on the system, simulations was performed for 3, 5, or 7 rounds, each consisting of 5 or 10 parallel trajectories of 40 ns. GROMACS was used to perform these FAST simulations, following the same main settings described in the system preparation section. The velocity-rescaling thermostat was applied to maintain the temperature at 310 K. Additional simulation details are provided in Ref.^36^.

For the CB1 and VP35 case studies (WT and mutants), we performed FAST adaptive sampling simulations to compare the WT and mutant systems and evaluate their effects on cryptic pocket dynamics. For VP35 and its mutants, we conducted 10 rounds of 10 parallel trajectories, each 50 ns long. For CB1, due to its larger system size, we performed 5 rounds of 5 parallel trajectories, each 40 ns long. For δ-secretase, to explore potential cryptic pockets, we ran 5 rounds of 10 parallel trajectories, each 40 ns in duration.

#### Molecular dynamics simulations

Some of the newly added proteins were simulated on the Folding@home distributed computing platform^60,61^ to collect ultra-long molecular dynamics trajectories. These simulations were performed using OpenMM^62^, with Amber03 force field and TIP3P explicit solvent. The main simulation settings were the same as those used in the FAST simulations, except that these runs were continuous (no rounds) and differed in trajectory lengths. For consistency, simulated structures included in the expanded training set were extracted from these trajectories using 40 ns windows, and the corresponding training labels were derived accordingly.

#### Sequence identity

To evaluate pairwise sequence similarity among proteins in the training dataset, we performed global pairwise sequence alignments using the BLOSUM62 substitution matrix with affine gap penalties (i.e., gap open = –11, gap extension = –1), consistent with BLASTP scoring conventions. Sequence identity was calculated as the fraction of identical residues in the aligned region, normalized by the length of the shorter sequence. A cutoff of 95% identity was applied to group proteins into clusters.

#### Cryptic pocket definition for analysis

For CB1 and δ-secretase, cryptic pocket residues were defined from holo structures as residues within 5 Å of the bound ligand, and pocket opening was monitored using geometric metrics derived from these ligand-contact residues. For VP35, we defined the cryptic pocket using prior combined experimental and computational characterization. Specifically, we analyzed C307 solvent exposure, quantified as SASA computed with the Shrake-Rupley algorithm^63^ implemented in MDTraj^64^ using a drug-sized probe (2.8 Å sphere), together with the G236-A306 distance. This distance can clearly separate open and closed pocket states.

### Network architectures

#### Geometric vector perceptron-based graph neural networks (GVP-GNNs)

AE-PocketMiner employs the same equivariant graph neural network as the original PocketMiner^36^, namely the geometric vector perceptron-based graph neural network (GVP-GNN). The GVP-GNN learns residue-level representations by capturing both the chemical and topological environments of residues through iterative information exchange with their neighboring residues.

The network takes as input both node features, which describe the intrinsic properties of each residue, and edge features, which encode the relationships between residue pairs. The node features include scalar values derived from the sine and cosine transformations of the backbone dihedral angles (φ, ψ, and ω), calculated using the atomic coordinates of C_i-1_, N_i_, C_i_, C_i_, and N_i+1_. In addition, the unit vector in the imputed direction of Cβ_i_ – Cα_i_ is computed by assuming tetrahedral geometry and normalizing the vector. To capture residue orientation, both forward and reverse unit vectors are included, along with a one-hot encoding representing the amino acid identity. The edge features are computed for the 30 nearest neighbor of each residue. These features include the unit vector pointing from Cα_j_ to Cα_i_, an encoding of the Euclidean distance ||Cα_j_ – Cα_i_||2 using Gaussian radial basis functions, and a sinusoidal encoding of the sequence separation (j – i) to represent the relative position along the backbone.

#### Attention module

To better capture local and long-range dependencies between residues, we incorporated a multi-head self-attention (MHA) mechanism with two heads, implemented in TensorFlow^65^. The attention module operates on the scalar features h_v output from the message passing layers. First, the scalar features are normalized using layer normalization to stabilize training and ensure consistent scaling across residues, producing normalized features h^-^_v. These normalized features are then passed to the MHA module, where the query (Q), key (K), and value (V) matrices are all derived from h^-^_v. This setup enables each residue to attend to every other residue (including itself) in the protein, allowing the model to capture both local and global structural dependencies. The attention-transformed features are subsequently combined with the original features via a residual connection to preserve the initial information while enriching it with context-aware representations. The combined representation is then fed into a feedforward network (FFN) followed by a sigmoid activation function, consistent with the original architecture, to produce the final cryptic pocket predictions (**Figure 1**).

In addition, we extract the attention scores, represented as a matrix produced by the MHA module. These scores are automatically normalized row-wise within the module, ensuring that the scores from each residue to all others (including itself) sum to one, which facilitates comparability across residues. The normalized scores can quantify residue-residue dependencies and provide interpretable insights into potential allosteric residues that may influence cryptic pocket formation, which is a key focus of this study.

#### Data featurization

As in our previous work^36^, we trained the model using protein structures sampled from FAST and conventional molecular dynamics simulations, incorporating both the original training dataset and the newly added simulations. To perform a case study on VP35, we removed all VP35 simulation data from the original training set to ensure the model was not biased by prior exposure.

For each simulated structure, every residue was assigned a binary label based on whether a pocket formed at that position within the following 40 ns of simulation. Consistent with our prior strategy, model training was conducted in two phases: 1.) using training labels derived from the LIGSITE pocket detection algorithm^59^, and 2.) using training labels derived from the fpocket algorithm^66^.

In the first phase, pockets were calculated using the LIGSITE pocket detection algorithm implemented in enspara^67^. The parameters used were: minimum rank = 7, grid spacing = 0.7 Å, probe radius = 1.4 Å, and minimum cluster size = 3 grid points. The detected pocket grid points were then mapped to nearby residues by assigning each pocket grid point to the nearest residue, following the same approach used in our previous work. For labeling, we computed the difference between each residue’s assigned pocket volume in the starting structure and its maximum pocket volume within the 40 ns simulation window. A residue was labeled as positive (1) if, at any point during the simulation (trajectories saved every 20 ps), its assigned pocket volume increases by at least 20 Å^3^ compared to the starting structure. Otherwise, it was labeled as negative (0).

In the second phase, we used the fpocket algorithm with default parameters to identify cavities (i.e., pockets) within each simulated protein structure. Unlike LIGSITE, fpocket also provides a druggability score for each pocket based on geometric and physicochemical features. To derive residue-level training labels, we iterated through all detected pockets and assigned each residue the maximum druggability score of any nearby pocket(s). Residues not belonging to any pocket were assigned a score of 0. Since fpockcet druggability score range from 0 to 1, these values were directly used as training labels without applying any thresholding.

#### Model training

We trained the model using the updated architecture describe above and the same set of hyperparameters from our previous work. Each residue node was represented with 8 vector and 50 scalar dimensions, while edges between nodes contained 1 vector and 32 scalar features. The hidden dimensions within the GVP layers were set to 16 vectors and 100 scalars. The model consisted of 4 encoding layers and used a neighbor list of 30 residues for graph propagation. A dropout rate of 0.1 was applied during training to prevent overfitting. All hyperparameters were originally proposed in the GVP-GNN publication^68^ and demonstrated little sensitivity in our prior evaluations, although smaller batch size yielded slightly better performance. In this study, we increased the batch size from 4 to 32 residues to accommodate a training set that was nearly three times bigger and to take advantage of the available GPU resources. The model was trained for 20 epochs using LIGSITE-derived labels, followed by 5 additional epochs using labels based on fpocket druggability scores. The final model was selected based on the highest validation AUC achieved across all training runs.

#### Model evaluation

After obtaining a single model, we evaluated the performance of AE-PocketMiner using the same test set from our previous work. The test set comprised 24 apo structures known to form ligand-binding cryptic pockets, 4 hyper-rigid proteins, and 7 proteins previously selected for extensive ligand screening. For details on how these proteins were selected, we refer readers to the original work^36^. All test proteins shared less than 45% sequence identity with those proteins in the training set, and those within the 40-45% range exhibited very limited structural homology.

### Experimental Measurements

#### Protein expression and purification

VP35 and its variant constructs were expressed in *E. coli* BL21(DE3) Gold cells (Agilent Technologies). Cells were grown in LB medium at 37 °C to an OD_600_ of ∼0.6–0.8 and induced with 1 mM IPTG (Gold Biotechnology), followed by a 24-hour expression at 18 °C. Cells were harvested by centrifugation and lysed by sonication in lysis buffer (20 mM sodium phosphate, 1 M NaCl, 5 mM β-mercaptoethanol, pH 8.0). Lysates were separated from insoluble materials by centrifugation and purified using Ni-NTA affinity chromatography, followed by TEV protease-mediated tag removal. Proteins were subsequently purified by cation-exchange chromatography and size-exclusion chromatography (Superdex 75). Final samples were maintained in 10 mM HEPES (pH 7.0), 150 mM NaCl, 1 mM MgCl₂, and 2 mM TCEP. Concentrations were determined from absorbance at 280 nm (ε = 6,970 M⁻¹cm⁻¹), and sample quality was verified by SDS-PAGE and intact mass analysis.

Purified human legumain (residues 18-323) was purchased from Biomatik in lyophilized condition.

#### Thiol labeling

Cysteine accessibility was quantified by labeling with 5,5′-dithiobis-(2-nitrobenzoic acid) (DTNB, Sigma). Labeling was initiated by mixing 10 µM protein with DTNB at varying concentrations in buffer (20 mM Tris, pH 8.0, 150 mM NaCl) at 25 °C. Absorbance at 412 nm, corresponding to TNB2-release, was recorded using a stopped-flow spectrophotometer (Applied Photophysics) until the signal plateaued (∼300 s). Raw kinetic traces were corrected by subtracting buffer-only controls and subsequently analyzed by global fitting to multi-exponential functions, with the number of exponentials corresponding to the number of cysteine residues in each construct. This approach generated the observed rate constants (6^$%^) for each site across DTNB concentrations. Data fitting was carried out in Python (SciPy), and values are reported as mean ± SD from at least three independent experiments. The dependence of 6^$%^ on DTNB concentration was analyzed using the Linderstrom-Lang model as previously described^69^. Depending on whether EXX or EX2 mechanism was followed, the equilibrium constant for pocket opening (Keq) was calculated.

## Data Availability

All data supporting the findings of this study are available within the article, the code repository, and the supplementary information files, or can be provided upon request.

## Code Availability

FAST and PocketMiner software packages are freely available on GitHub at https://github.com/bowman-lab/fast and https://github.com/Mickdub/gvp.

## Acknowledgements

We gratefully acknowledge the citizen scientists who participate in Folding@home for generously contributing their personal computers to run simulations and generate data. This work was funded by the National Institutes of Health through NIGMS R35GM152085, and NSF MCB 2218156. We thank Dr. Artur Meller for valuable discussions and insightful advice on model training.

## Author Contributions

S.Z. and G.R.B. conceptualized the project direction and strategy. S.Z. and R.K. collaboratively explored and tested model architectures to achieve the project objectives. S.Z. processed the simulation data, generated training labels, and performed model training and evaluation. P.M. and D.K. conducted experiments and data analysis. S.Z. drafted the manuscript, with editorial contributions from P.M., D.K. and R.K. G.R.B. proposed the initial project idea, provided edits to the manuscript, and acquired funding.

